# Viral overexpression of human alpha-synuclein in mouse substantia nigra dopamine neurons results in hyperdopaminergia but no neurodegeneration

**DOI:** 10.1101/2024.05.03.592188

**Authors:** Sofia Ines Garcia Moreno, Fabian Limani, Iina Ludwig, Catherine Gilbert, Christian Pifl, Thomas S. Hnasko, Thomas Steinkellner

## Abstract

Loss of select neuronal populations such as midbrain dopamine (DA) neurons is a pathological hallmark of Parkinson’s disease (PD). The small neuronal protein α-synuclein has been related both genetically and neuropathologically to PD, yet how it contributes to selective vulnerability remains elusive. Here, we describe the generation of a novel adeno-associated viral vector (AAV) for Cre-dependent overexpression of wild-type human α-synuclein. Our strategy allows us to restrict α-synuclein to select neuronal populations and hence investigate the cell-autonomous effects of elevated α-synuclein in genetically-defined cell types. Since DA neurons in the substantia nigra *pars compacta* (SNc) are particularly vulnerable in PD, we investigated in more detail the effects of increased α-synuclein in these cells. AAV-mediated overexpression of wildtype human α-synuclein in SNc DA neurons increased the levels of α-synuclein within these cells and augmented phosphorylation of α-synuclein at serine-129, which is considered a pathological feature of PD and other synucleinopathies. However, despite abundant α-synuclein overexpression and hyperphosphorylation we did not observe any DA neurodegeneration up to 90 days post virus infusion. In contrast, we noticed that overexpression of α-synuclein resulted in increased locomotor activity and elevated striatal DA levels suggesting that α-synuclein enhanced dopaminergic activity. We therefore conclude that cell-autonomous effects of elevated α-synuclein are not sufficient to trigger acute DA neurodegeneration.

## Introduction

Alpha-synuclein is a small protein of 140 amino acids that is closely related to the pathogenesis of Parkinson’s disease (PD) since it was discovered that a mutation in the α-synuclein gene (SNCA) causes a familial form of the disease [37]. Around the same time, α-synuclein was shown to be a major protein constituent of Lewy bodies (LB) in postmortem brains of patients with PD and dementia with LBs (DLB) [42]. Later, other mutations in the SNCA gene suspected to cause familial autosomal dominant PD (*e.g.* A30P) were described including triplication and duplication of the wildtype SNCA gene [5, 25, 41, 47]. The genetic findings seemed to imply that mutated forms of α-synuclein or elevated levels of the wildtype protein are toxic and capable of inducing neurodegeneration.

It remains, however, unclear how gene mutations or multiplications of SNCA would lead to selective degeneration of specific neuronal populations despite widespread expression of α-synuclein in neurons and many other cell types [7, 12, 18, 34]. PD is largely characterized by intraneuronal LBs throughout the brain, but the cardinal motor symptoms are largely caused by profound degeneration of dopamine (DA) neurons in the substantia nigra *pars compacta* (SNc) [8, 16]. The degeneration of specific cholinergic, noradrenergic, serotonergic and other neuronal populations has also been described but the cell loss in these structures is less well defined [13].

To test the ‘gain-of-toxic-function’ hypothesis of α-synuclein, numerous animal models in which α-synuclein levels are elevated have been generated.

For instance, transgenic mice in which α-synuclein expression is driven by either ubiquitous, neuronal or more specific promoters have been created (for summary see [6]). While some of these lines develop extensive α-synuclein expression and LB-like inclusions throughout the brain they usually do not display the selective cell loss observed in the human disease, *i.e.* degeneration of DA neurons in the SNc [6].

Other animal models have been established in which expression of α-synuclein was more restricted to vulnerable regions of interest through stereotaxic injections of viral vectors such as adeno-associated viruses (AAVs) or lentiviruses [22, 24, 29]. Viral vectors have the advantage that they can be delivered to adult animals of different ages and into brain regions of interest including those most relevant to the disease such as the SNc in the ventral midbrain. Many of these models lead to moderate DA neurodegeneration, but the vectors to deliver α-synuclein do not cause cell-type specific expression. Hence, infusion of viral particles into *e.g.* the ventral midbrain induces overexpression of α-synuclein in many different cell types including DA and non-DA neurons. Therefore, a conclusion about whether elevation of α-synuclein within DA neurons suffices to trigger degeneration remains unclear.

We therefore sought to generate a novel Cre recombinase-dependent AAV to allow cell-type specific expression of α-synuclein in defined neuronal populations. To determine in more detail whether cell-autonomous overexpression of α-synuclein would be sufficient to trigger DA neurodegeneration, we restricted human wildtype α-synuclein to SNc DA neurons of DAT^Cre^ mice. Despite abundant overexpression of α-synuclein in DA neurons and increased phosphorylation at serine-129 we did not observe any appreciable degeneration of DA neuron terminals in the striatum or loss of cell bodies in the SNc up to 90 days after virus infusion.

In contrast, overexpression of α-synuclein in SNc DA neurons led to a hyperdopaminergic phenotype characterized by elevated locomotor activity and increased striatal DA tissue levels. Together, our results indicate that cell-autonomous overexpression of α-synuclein in DA neurons can drive DA dysfunction but is not sufficient to induce acute DA neurodegeneration.

## Materials and Methods

### Animals

We used adult mice expressing Cre recombinase under the control of the dopamine transporter (DAT^Cre^: B6.SJL-Slc6a3tm1.1(cre)Bkmn/J, Jackson stock 006660), choline-acetyl transferase promoter (ChAT^Cre^: B6;129S6-Chattm2(cre)Lowl/J, Jackson stock 006410), adenosine A2a receptor (A2a^Cre^; Tg(Adora2a-cre)K139Gsat/Mmcd, Jackson stock 036158) from The Jackson Laboratory, Bar Harbor, ME. Dopamine receptor D_1_ ^Cre^ mice (Drd1a^+/Cre^) have been obtained from the lab of Richard Palmiter [15]. All mice were 12-20 weeks of age at the time of stereotaxic surgery and were bred on a C57BL/6J genetic background. Mice were group-housed, and maintained on a 12:12-hour light:dark cycle with food and water available ad libitum.

### Generation of Cre-dependent adeno-associated viruses

A plasmid containing cDNA for human α-synuclein wildtype (in a pLV backbone) was a gift from Brian Spencer and Robert Rissman (UCSD). Human α-synuclein cDNA was amplified by PCR and AscI restriction sites added at 5’ and 3’ ends for insertion into pAAV-FLEX-hSyn1 to generate a Cre-dependent vector. DNA sequence was verified by Sanger sequencing and tested for Cre-mediated recombination and expression in HEK-293 cells. DNA was packaged into AAV_DJ_ serotype by Vigene Biosciences (now Charles River Laboratories). The construct will be made available on Addgene.

### Stereotaxic surgeries and viral infusions

Mice were anaesthetized with isoflurane (1-4%) and placed into a stereotaxic frame (RWD Life Science Co., Guangdong, China). Replication-incompetent AAV_DJ_ serotype was used to drive expression of eGFP or human α-synuclein under the control of the human Synapsin 1 promoter: AAV_DJ_-hSyn1-DIO-eGFP (3.6×10^13^ genome copies per ml [gc/ml]), AAV_DJ_-hSyn1-DIO-hASYN-WT (1.5×10^13^ gc/ml). The GFP virus was packaged at the Salk GT3 vector core (La Jolla, CA). Viruses were microinfused (300 nL) into the left SNc (−3.4 AP, −1.25 ML, −4.25 DV) using a 30G stainless steel injectors at 100 nL/min.

### Behavior

*Open-field*: Horizontal locomotion (total distance traveled) was measured in square boxes (36 cm x 36 cm x 45 cm) using a video camera mounted above the box and analyzed using ANY-maze software (Stoelting). Distances traveled were recorded for 30 min. Mice were tested either at 21 and/or 90 days after virus infusion.

### Rotarod

Motor coordination was measured using an automated rotarod system for rodents (MedAssociates). Mice were placed onto a rotating drum (accelerating speed from 4 to 40 revolutions per minute over 300 sec) for three trials separated by a 15 min inter-trial interval. No training period was done prior to the test phase. The latency to fall off was measured by integrated laser beam detectors. Mice were tested at 90 days after virus infusion.

### Histology

Mice were deeply anaesthetized with pentobarbital (100 mg/kg i.p.; Exagon^®^ 500 mg/mL, Richter Pharma) and transcardially perfused with 10–20 mL of phosphate-buffered saline (PBS) 21 or 90 days after viral injection. This was followed by perfusion with 60-70 mL of 4% paraformaldehyde (PFA) at a rate of ca. 6 ml/min. Brains were extracted, post-fixed in 4% PFA at 4°C overnight, and cryoprotected in 30% sucrose in PBS for 48–72 hours at 4°C. Brains were snap-frozen on dry ice or in liquid nitrogen and stored at −80°C. Coronal sections (30 μm) were cut using a cryostat (CM3050S, Leica, Wetzlar, Germany) and collected in PBS containing 0.01% sodium azide.

For fluorescent immunostaining, brain sections were blocked with 5% normal donkey serum in PBS containing 0.3% Triton X-100 (blocking buffer) (1 hour, room temperature). Sections were then incubated with one or more of the following primary antibodies in blocking buffer overnight at 4°C (rabbit anti-TH, 1:2000; AB152, Merck-Millipore; sheep anti-TH, 1:2000, P60101-150, Pel Freez; rat anti-human α-synuclein, 15G7, Enzo Life Sciences; mouse anti-α-synuclein [Syn1, clone 4D6], 1:1000, 834301, BioLegend; mouse anti-phospho-serine-129 α-synuclein, 1:1000, 825701 [formerly Covance MMS-5091], BioLegend).

Sections were rinsed 3 x 15 min with PBS and incubated in appropriate secondary antibodies (Jackson ImmunoResearch, West Grove, PA) conjugated to Alexa 488, Alexa 594 or Alexa 647 fluorescent dyes (5 μg/ml) (2 hours, room temperature). Sections were washed 3 x 15 min with PBS, mounted onto glass slides and coverslipped with Fluoromount-G mounting medium (Southern Biotech, Birmingham, AL) supplemented with DAPI (0.5 µg/ml, Roche, Basel, Switzerland). Images were acquired using either a Zeiss AxioObserver epifluorescence microscope with Apotome, TissueFAXS PLUS or Nikon Eclipse TS100 confocal microscope.

For the chromogenic staining, free-floating sections (30 µm) were washed 3 times (5 minutes) in 0.1 M Tris-buffered saline, pH = 7.6 (TBS), before incubation of sections in 3% H_2_O_2_ (in TBS) for 30 minutes at room temperature to quench endogenous peroxidases followed by blocking in 5% normal donkey serum/0.3% Triton X-100 in TBS for 1 hour at room temperature. Rabbit anti-TH (ab152) was used at a concentration of 1:2000 in blocking buffer. Sections were incubated in primary antibody solutions overnight at 4°C. The following day, sections were washed 3 times (15 minutes) in TBS and incubated with a donkey antirabbit biotinylated secondary antibody (Jackson ImmunoResearch Laboratories) at 1:400 in blocking buffer for 2 hours at room temperature. Sections were again washed 3 times (15 minutes) in TBS and incubated in avidin-biotin complex solution (Vectastain Elite ABC kit, Vector Laboratories) for 2 hours at room temperature before additional washes in TBS (2 times, 10 minutes). Sections were incubated in DAB solution (0.4 mg/mL 3,3-diaminobenzidine–HCl and 0.005% H2O2 in 0.1 M Tris-HCl) for 3–5 minutes at room temperature. Sections were again rinsed 2 times in TBS before mounting onto glass slides and dried overnight. The next day, sections were dehydrated through increasing concentrations of ethanol, cleared with CitriSolv (Thermo Fisher Scientific), and cover-slipped using VectaMount® Mounting Media.

### Stereology

Stereological sampling was performed using Stereo Investigator software (MBF Europe B.V.) by an investigator blind to treatment as described [44]. Counting frames (100 × 100 μm) were randomly placed on a counting grid (200 × 200 μm) and sampled using a 7 μm optical dissector with guard zones of 10% of the total slice thickness on each site (∼2 μm). The boundaries of the SNc were outlined under magnification (4x objective). Cells were counted with a 20x objective (0.45 numerical aperture) using an Olympus BX51 microscope. A dopaminergic neuron was defined as an in-focus TH–immunoreactive (TH-IR) cell body with a TH-negative nucleus within the counting frame. Every fifth section was processed for TH-IR, resulting in 6 sections sampled per mouse, and every section was counted. The number of neurons in the SNc was estimated using the optical fractionator method, which is unaffected by changes in the volume of reference of the structure sampled. Between 70 and 160 objects per animal were counted to generate the stereological estimates.

### Densitometry

TH^+^ fibers in the striatum were imaged using a TissueFAXS PLUS fluorescence slide scanner with a 20x objective. The average intensity of TH staining in a delineated region of the striatum was quantified with ImageJ software (NIH). The relative optical density of TH^+^ fibers was normalized by subtracting the background intensity of the cortex and using the particle analysis function. Fluorescence images of Syn1 or p-syn stained sections in in the midbrain were captured with a Nikon Eclipse TS100 confocal microscope. The intensity of Syn1 or p-syn fluorescence was analyzed by subtracting background intensity and using the particle analysis function in ImageJ.

### Neurochemistry

Mice were anesthetized with pentobarbital as described above and killed by decapitation, their brains isolated and immediately frozen in liquid nitrogen and stored at −80°C. Left and right striata were dissected in a frozen state and weighed to obtain wet weights. 50 volumes of 0.1 M perchloric acid containing 0.4 mM NaHSO_3_ were added to striata for homogenization using a tip sonicator. Homogenates were then centrifuged for 10 min at 4°C at 14,000 x rpm. To measure dopamine and the metabolites 3,4-dihydroxyphenylacetic acid and homovanillic acid, we injected 100 µL of supernatants directly into a high-performance liquid chromatography (HPLC) system equipped with a LiChroCART^®^ 250-4, RP-18, 5 μm (VWR) column and a BAS (Bioanalytical System) electrochemical detector. The mobile phase consisted of 0.1 M sodium acetate buffer, pH 4.3, containing 1 mM EDTA, 0.2 mM l-heptane sulfonic acid, 0.1% triethylamine, 0.2% tetrahydrofuran and 10% methanol. Purchased standards (Sigma-Aldrich) for all detected compounds were utilized for quantification.

### Immunoblotting

Tissue punches from the striatum were prepared using a disposable biopsy puncher (Integra, #3331-A) from fresh-frozen brains of unilaterally injected mice in a cryostat and kept at −80°C until further processing. Proteins were extracted in ice-cold lysis buffer containing 150 mM NaCl, 50 mM Tris–HCl, and 1% Triton X-100, pH 7.4 and supplemented with protease inhibitors (Sigma, Complete Mini Protease Inhibitor Cocktail, #11836153001) using a probe tip sonicator (Sonics, Vibra Cell). Homogenates were further incubated on a rotator for 1 hr at 4°C and centrifuged at 12,000 x g for 15 min at4°C. Supernatant was kept, and protein concentration was measured using the BCA method (Pierce BCA Protein Assay, Thermo Fisher, #23225).

Samples were incubated in 2x Laemmli Buffer (120 mM Tris–HCl, pH = 6.8, 20% [w/v] glycerol, 4% sodium dodecyl sulfate, 10% 2-mercaptoethanol, 0.02% bromphenol blue) for 5 min at 95°C, and 10 µg of protein was separated on a 4–20% sodium dodecyl sulfate polyacrylamide gel electrophoresis (Bio-Rad, #4561096) and blotted onto nitrocellulose membranes by wet transfer. Membranes were fixed in 0.4% PFA in PBS for 30 min at room temperature (to increase binding of monomeric α-synuclein to the membrane and stained with Ponceau S solution to determine protein loading. Blocking was performed in 0.1% PBST (phosphate-buffered saline [PBS], 0.1% Tween-20) containing 5% non-fat dry milk for 1 hr at room temperature. Incubation with primary antibodies (rat anti-human α-synuclein, 15G7, 1:100 Enzo Life Sciences; mouse anti-α-synuclein [Syn1, clone 4D6], 1:1000, 834301, BioLegend; rabbit anti-GAPDH 1:2000 (D16H11) Cell Signaling Technologies) in blocking buffer was done overnight at 4°C. Next day, membranes were washed three times in 0.1% PBST and incubated with secondary antibodies (donkey anti-mouse-Alexa-488, donkey Anti-rat-Alexa-594 and donkey Anti-rabbit-Alexa-647; 1:1000; Jackson Immuno Research) in blocking solution for 2 hr at room temperature before another three 15 min washes in 0.1% PBST. Membranes were briefly rinsed in PBS before imaging using the Bio-Rad ChemiDoc MP imaging system. Band densities were quantified using Image J (NIH), normalized to GAPDH (loading control; ca. 37 kD), and are expressed as relative intensities.

### Statistics

GraphPad Prism (GraphPad Software Inc., San Diego, CA) was used to analyze data and create graphs. All data are expressed as means ± SEM. Data were analyzed by two-tailed unpaired Student’s t-test, one-or two-way ANOVA followed by post-hoc multiple comparisons tests as indicated in the respective figure legends.

## Results

### Cell-type specific expression of α-synuclein in different neuronal populations

We generated a novel viral vector for cell-type specific expression of human wildtype α-synuclein. Therefore, we cloned wildtype human α-synuclein into a double-inverted open reading frame plasmid (pAAV-hSyn1-DIO). The orientation of α-synuclein in this vector is inverted and flanked by loxP and lox2272 sites (**Fig. 1A**). Upon recognition by Cre recombinase the transgene is flipped, locked in the correct open reading frame and can be expressed [40].

**Figure 1:**
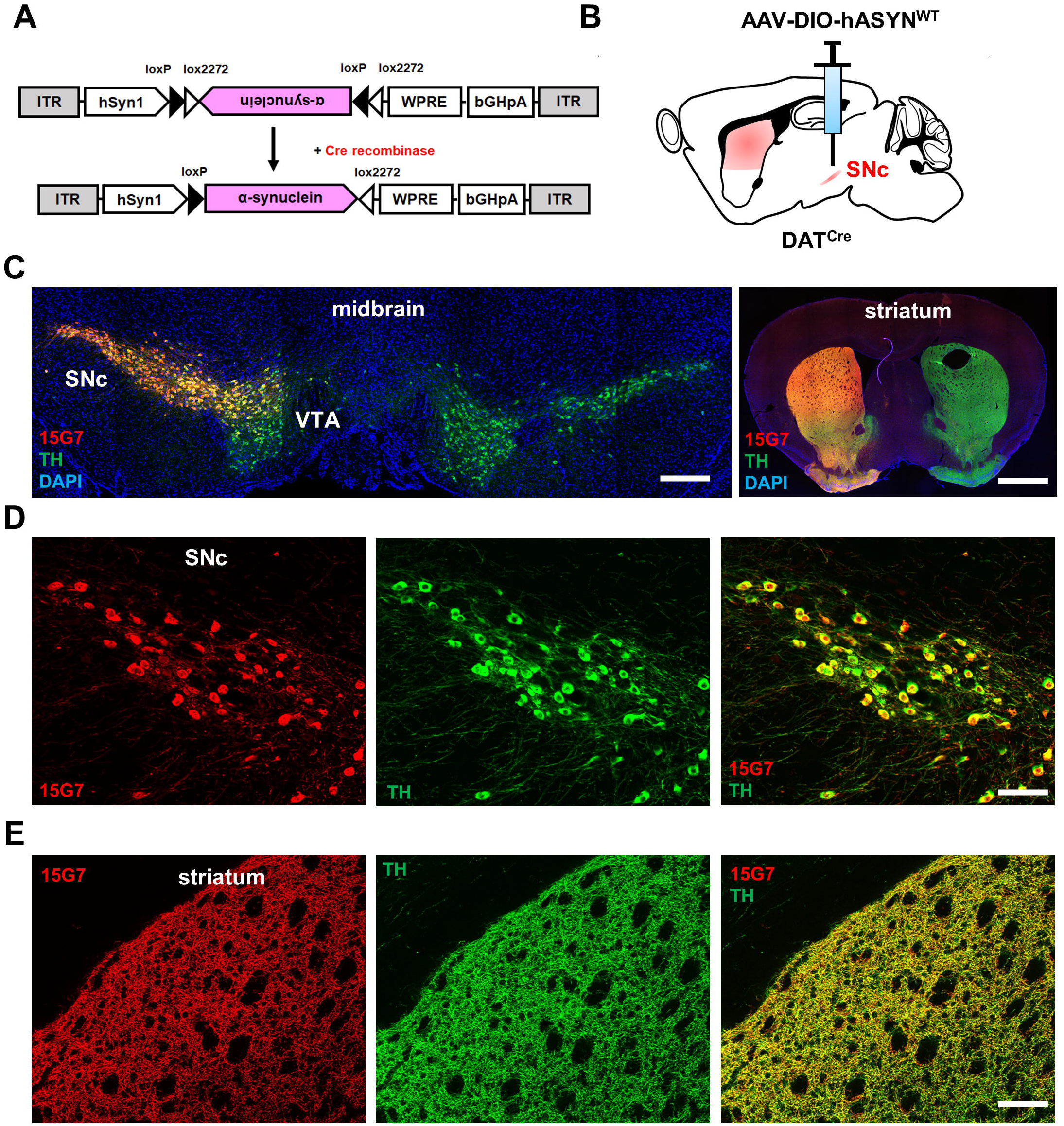
Cell-type specific expression of human alpha-synuclein in mouse dopamine neurons. **(A)** AAV construct showing that in the presence of Cre recombinase, α-synuclein is recombined to allow for cell-type-specific expression. **(B)** Strategy for unilateral expression of AAV-DIO-hASYN^WT^ in the SNc of DAT^Cre^ mice. **(C)** Immunohistochemistry shows expression of human α-synuclein (red), detected in TH+ DA neurons (green) in coronal sections through SNc (left panel) and striatum (right panel) using the human α-synuclein-specific antibody 15G7. Counterstaining was done with DAPI (blue). Scale bar: 500 µm. **(D, E)** Higher magnification images show co-expression of human α-synuclein (red) with the dopamine marker TH (green) in SNc neurons (scale bar: 100 µm) **(D)** and striatal terminals (scale bar: 200 µm) **(E)**.

To test Cre-dependence and expression *in vivo*, the construct was packaged into an adeno-associated virus (AAV_DJ_) and stereotaxically injected into different Cre driver lines or non-Cre expressing wildtype mice (**Fig. 1B-E** and **Supplemental Fig. 1**). Twenty-one days after infusion of AAV-DIO-hASYN^WT^ into the left substantia nigra of mice expressing Cre in dopamine neurons (DAT^Cre^ mice), we performed histology and stained midbrain and striatal sections for human α-synuclein using a human isoform-specific antibody (15G7) and an antibody against tyrosine hydroxylase (TH) (**Fig. 1C-E**). Injection of AAV-DIO-hASYN^WT^ into DAT^Cre^ mice demonstrates that expression was restricted to TH^+^ dopamine neurons in striatal terminals and in nigral cell bodies: out of 791 human α-synuclein (15G7)-positive cells counted from three different mice 791 cells were also TH-positive.

Similarly, human α-synuclein was only detectable with 15G7 in choline acetyltransferase (ChAT)-positive striatal cholinergic interneurons after injection of AAV-DIO-hASYN^WT^ into the dorsal caudate/putamen (CPu) of ChAT^Cre^ mice: out of 70 15G7^+^ cells counted (n=2 mice), 70 were ChAT^+^ (**Supplemental Fig. 1B**); and 15G7 staining was restricted to either the direct (hASYN expression striatal projections to the substantia nigra pars reticulata [SNr] and globus pallidus internus [GPi]) or indirect (hASYN expression in striatal projections to globus pallidus externus [GPe]) pathway after expression in D_1_ ^Cre^ or A ^Cre^ mice, respectively, demonstrated by co-labeling with either dopamine D_1_ or D_2_ receptors (**Supplemental Fig. 1C-D**).

Importantly, no α-synuclein was detected using the 15G7 antibody when AAV-DIO-hASYN^WT^ was injected into the midbrain of C57BL6/j wildtype mice confirming that there was no leaky transgene expression (**Supplemental Fig. 1A**).

### Quantification of α-synuclein levels in dopamine neurons after viral overexpression

To estimate whether levels of α-synuclein after virus overexpression within the dopamine system of DAT^Cre^ mice were indeed higher than endogenous levels, we stained midbrain sections with an α-synuclein antibody (Syn1) which recognizes both mouse and human α-synuclein (**Fig. 2A-C**). We then imaged injected (ipsilateral) and uninjected (contralateral) midbrain dopamine neurons in the substantia nigra by confocal microscopy (**Fig. 2B-C**) to quantify α-synuclein intensities. Syn1 staining was significantly higher on the injected (ipsilateral) compared to the non-injected (contralateral) side (**Fig. 2D**). Note that endogenous α-synuclein is usually not well detectable at the cell body level, but mostly localizes to presynaptic terminals unless it is overexpressed [32, 45].

**Figure 2:**
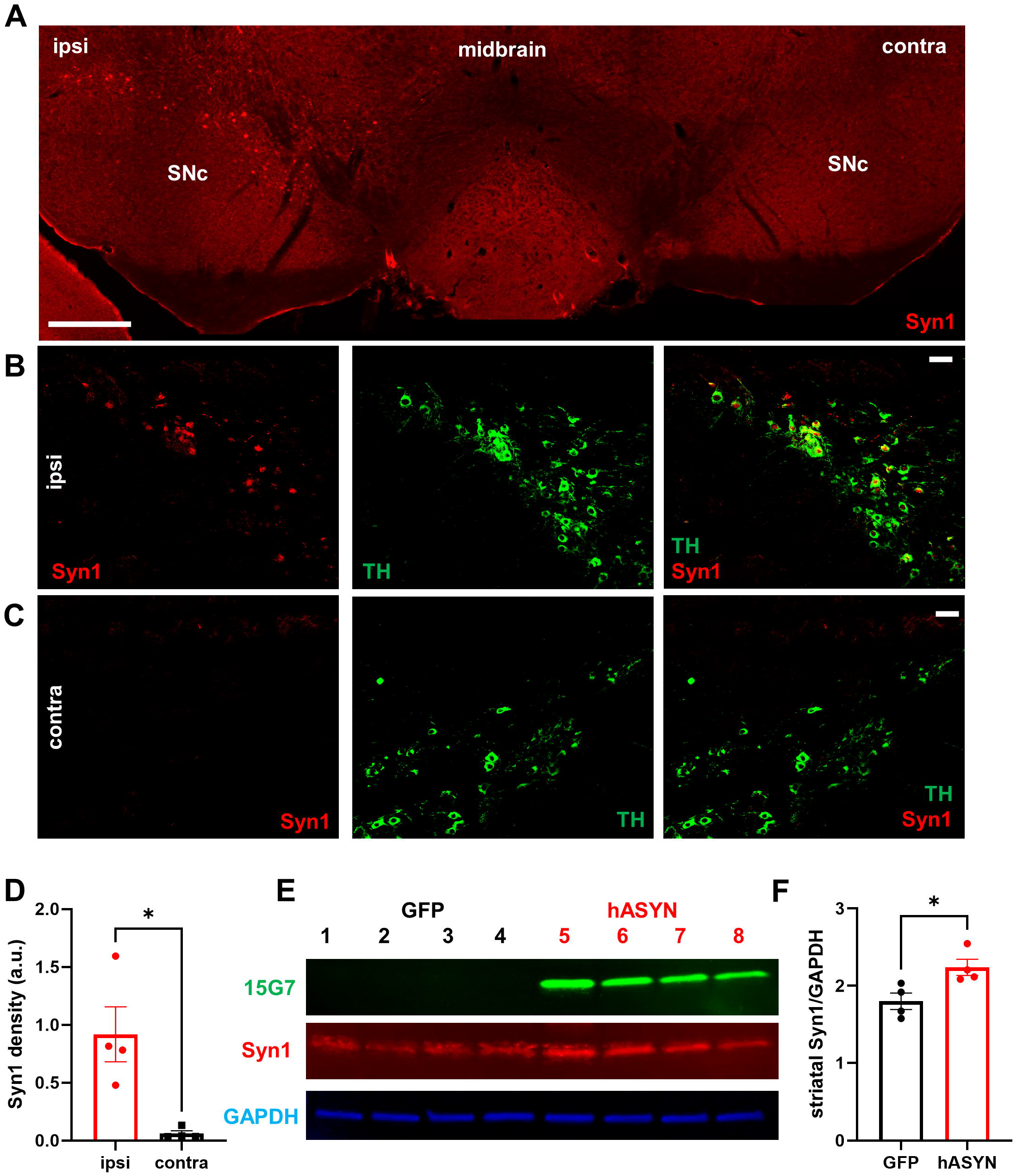
Overexpression of human alpha-synuclein leads to overall increase of alpha-synuclein in dopamine neurons. **(A)** Detection of total (endogenous mouse + overexpressed human) α-synuclein levels using the Syn1 antibody (red) in a midbrain section from a mouse unilaterally overexpressing human α-synuclein 21 days post infusion (dpi); scale bar: 500 µm. **(B-C)** Higher resolution confocal images of SNc neurons stained for Syn1 (red) and TH (green) on the ipsilateral **(B)** or contralateral **(C)** side 21 dpi; scale bars: 50 µm. **(D)** Densitometric quantification of total alpha-synuclein levels in midbrain DA neurons. n=4 mice/group; two-tailed paired Student’s t-test; t=3.395, df=3, *p=0.0426. **(E)** Immunoblot of striatal protein lysates probed for human α-synuclein (15G7; green), total α-synuclein (Syn1; red) and loading control (GAPDH; blue) n=4 mice/group. **(F)** Densitometric quantification of Syn1 levels normalized to GAPDH; n=4 mice/group; two-tailed unpaired Student’s t-test; t=2.937, df=6, * p=0.0260.

Additionally, we prepared striatal tissue punches for protein extraction to detect α-synuclein by immunoblotting. Staining with the human-specific antibody (15G7) confirms that human α-synuclein is only expressed after AAV-DIO-hASYN^WT^ but not AAV-DIO-eGFP injection (**Fig. 2E**). Immunoblotting with the Syn1 antibody suggested that total α-synuclein protein was again significantly increased after AAV-DIO-hASYN^WT^ overexpression compared to AAV-DIO-eGFP (**Fig. 2E-F**). However, it is important to mention that quantifications by both histochemistry and immunoblotting are biased because endogenous mouse α-synuclein cannot be easily subtracted in such an analysis and hence precise absolute quantification is difficult.

### Overexpression of human α-synuclein in dopamine neurons increases motor activity

To determine whether AAV-DIO-hASYN^WT^ would alter motor activity we conducted several behavioral tests. Locomotion was recorded in the open field test 21 and 90 days after unilateral virus infusion into the left substantia nigra of DAT^Cre^ mice. Surprisingly, mice injected with AAV-DIO-hASYN^WT^ in dopamine neurons displayed significantly increased motor activity during a 30 minute session both 21 and 90 days after injection compared to mice injected with AAV-DIO-eGFP (**Fig. 3A-D**). However, mice did not display any significant differences in rotational behavior at either time-point (**Fig. 3E**). Similarly, performance in the accelerating rotarod (latency to fall off) was not significantly different between α-synuclein and GFP mice 90 days after virus infusions (**Fig. 3G**).

**Figure 3:**
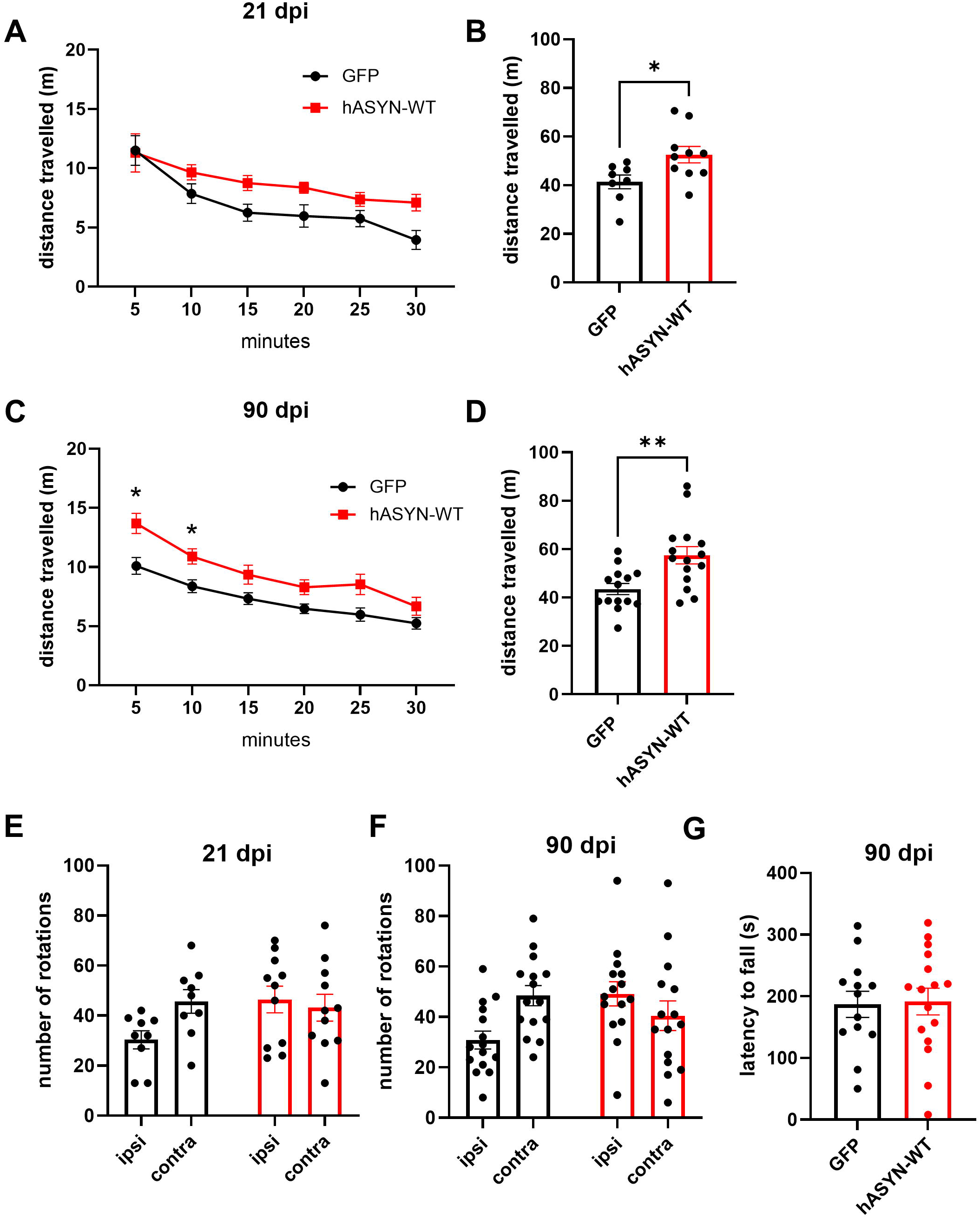
Mice with unilateral alpha-synuclein overexpression in dopamine neurons display increased locomotor activity. **(A-D)** DAT^Cre^ mice were unilaterally injected with AAV-DIO-hASYN^WT^ and tested in an open-field arena either 21 **(A, B)** or 90 **(C, D)** days after surgery. Left panels **(A, C)** show time courses of 5-min bins over 30 minutes (two-way ANOVA followed by Tukey’s multiple comparisons test). Right panels **(B, D)** show cumulative distances traveled over 30 minutes (*21 days*: n=8 GFP mice and n=10 hASYN mice; two-tailed unpaired Student’s t-test: t=2.478, df=16, *p=0.0248; *90 days*: n=14 GFP mice and n=15 hASYN mice; two-tailed unpaired Student’s t-test: t=3.246, df=27, **p=0.0031). **(E-F)** Number of ipsi- and contraversive rotations is unchanged hASYN and control (GFP) overexpressing mice, both 21 **(E)** and 90 days **(F)** after viral infusion. **(G)** Rotarod test did not reveal differences in the latency to fall in AAV-DIO-hASYN^WT^ or GFP mice 90 days after injection.

### Increased striatal dopamine levels but unchanged dopamine neuron number in mice after α-synuclein overexpression

Next, we analyzed whether unilateral infusion of AAV-DIO-hASYN^WT^ into the SNc of DAT^Cre^ mice would alter dopamine levels in the striatum. We isolated striatal tissue punches from mice injected with either AAV-DIO-hASYN^WT^ or AAV-DIO-eGFP 90 days after infusion and measured dopamine and its major metabolites by HPLC with electrochemical detection. Striatal dopamine tissue levels were significantly increased in mice overexpressing human α-synuclein compared to GFP controls (**Fig. 4A**), while the dopamine metabolites 3,4-dihydroxyphenylacetic acid (DOPAC) and homovanillic acid (HVA) remained largely unchanged (**Fig. 4B-C**). The molar ratios of DOPAC/dopamine or HVA/dopamine were unaltered (**Supplemental Fig. 3**).

**Figure 4:**
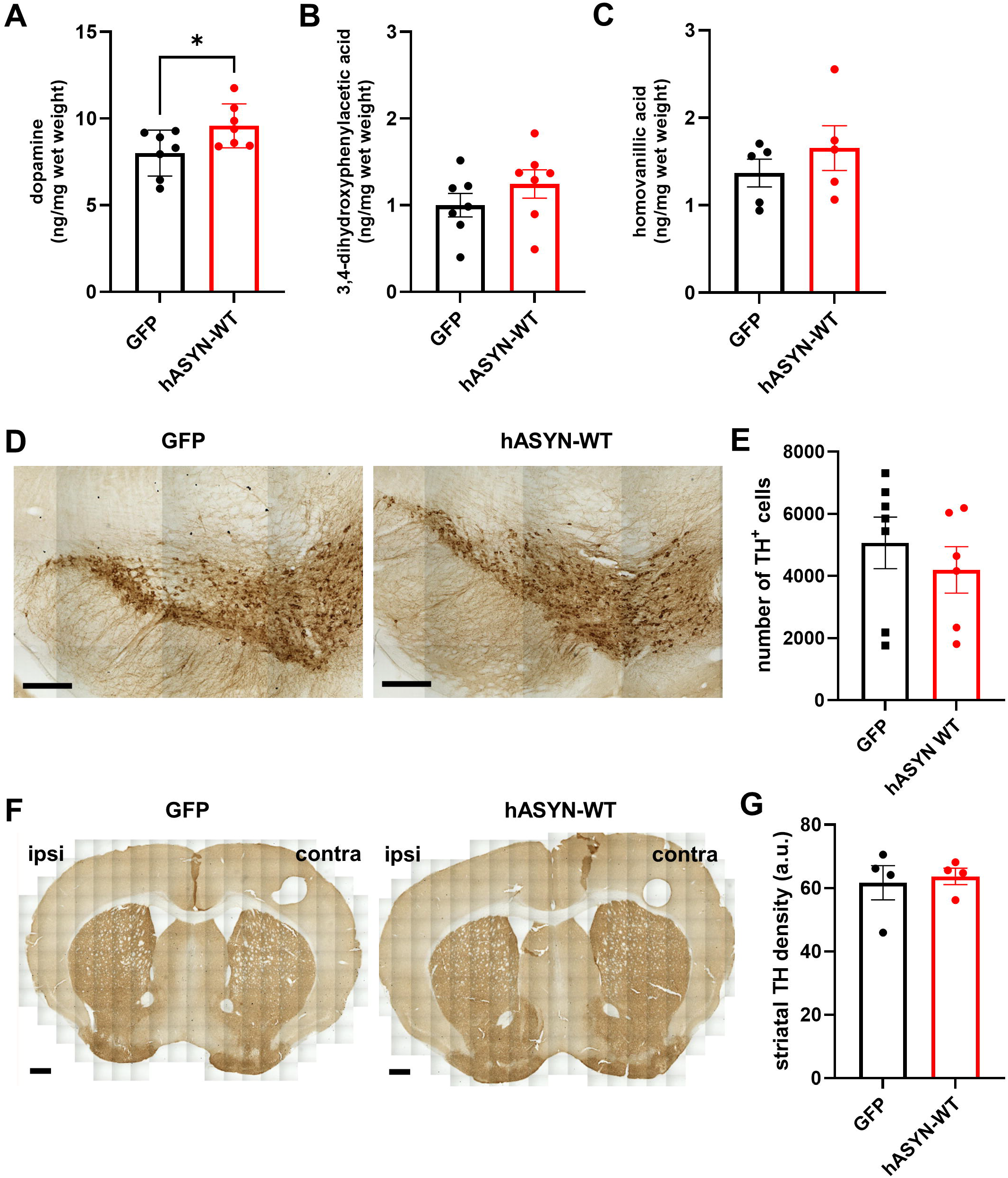
Overexpression of alpha-synuclein in dopamine neurons increases striatal dopamine levels without altering the total number of TH neurons. **(A-C)** Striata from mice overexpressing AAV-DIO-hASYN^WT^ or GFP (control) were extracted and dopamine **(A)**, 3,4-dihydroxyphenylacetic acid **(B)** and homovanillic acid **(C)** were measured by high-performance liquid chromatography with electrochemical detection. n=7/group for DA (two-tailed unpaired Student’s t-test: t=2.263, df=12, *p=0.0429) and DOPAC and n=5/group for HVA measurements. **(D)** Representative midbrain sections from GFP (left) and AAV-DIO-hASYN^WT^ (right) mice stained for TH 90 days after unilateral virus infusion. Scale bar: 200 µm. **(E)** Number of TH^+^ cells were counted in the left SNc by unbiased stereology; n=7 for GFP group and n=6 for hASYN group. **(F)** Representative striatal sections of GFP (left) and AAV-DIO-hASYN^WT^ (right) stained for TH-DAB. Scale bar: 500 µm. **(G)** Densitometric analysis of ipsilateral (left) striata; n=4/group.

To determine whether α-synuclein overexpression would affect the number of dopamine neurons in the substantia nigra, we counted TH^+^ cells on the side of injection using unbiased stereological cell counting. There were no significant differences in the estimated total number of TH^+^ neurons between α-synuclein overexpressing or GFP control mice 90 days after virus infusion (**Fig. 4D-E**). Similarly, striatal TH density in the dorsal striatum remained unchanged on the ipsilateral sides between α-synuclein and GFP mice at 90 days post injection (**Fig. 4F-G**).

### Increased serine-129 phosphorylation of α-synuclein after overexpression

Phosphorylation of α-synuclein at serine-129 is considered a pathological hallmark of synucleinopathies [3, 10] but has more recently been shown to also facilitate interaction with protein partners and to serve as a marker for synaptic activity [35, 38]. To determine whether the increase in locomotor activity seen after α-synuclein overexpression and the increased striatal tissue levels in these mice would be accompanied by altered α-synuclein serine-129 phosphorylation, we stained midbrain sections from AAV-DIO-hASYN^WT^ injected DAT^Cre^ mice with a phospho-serine-129 specific antibody at 21 and 90 days post virus infusion (**Fig. 5A-C**). We found that phospho-serine-129 staining was significantly higher on the side of α-synuclein overexpression at both time-points, while the contralateral sides showed only background staining (**Fig. 5D-E**) suggesting that activity of dopamine neurons on the side of α-synuclein overexpression may have increased.

**Figure 5:**
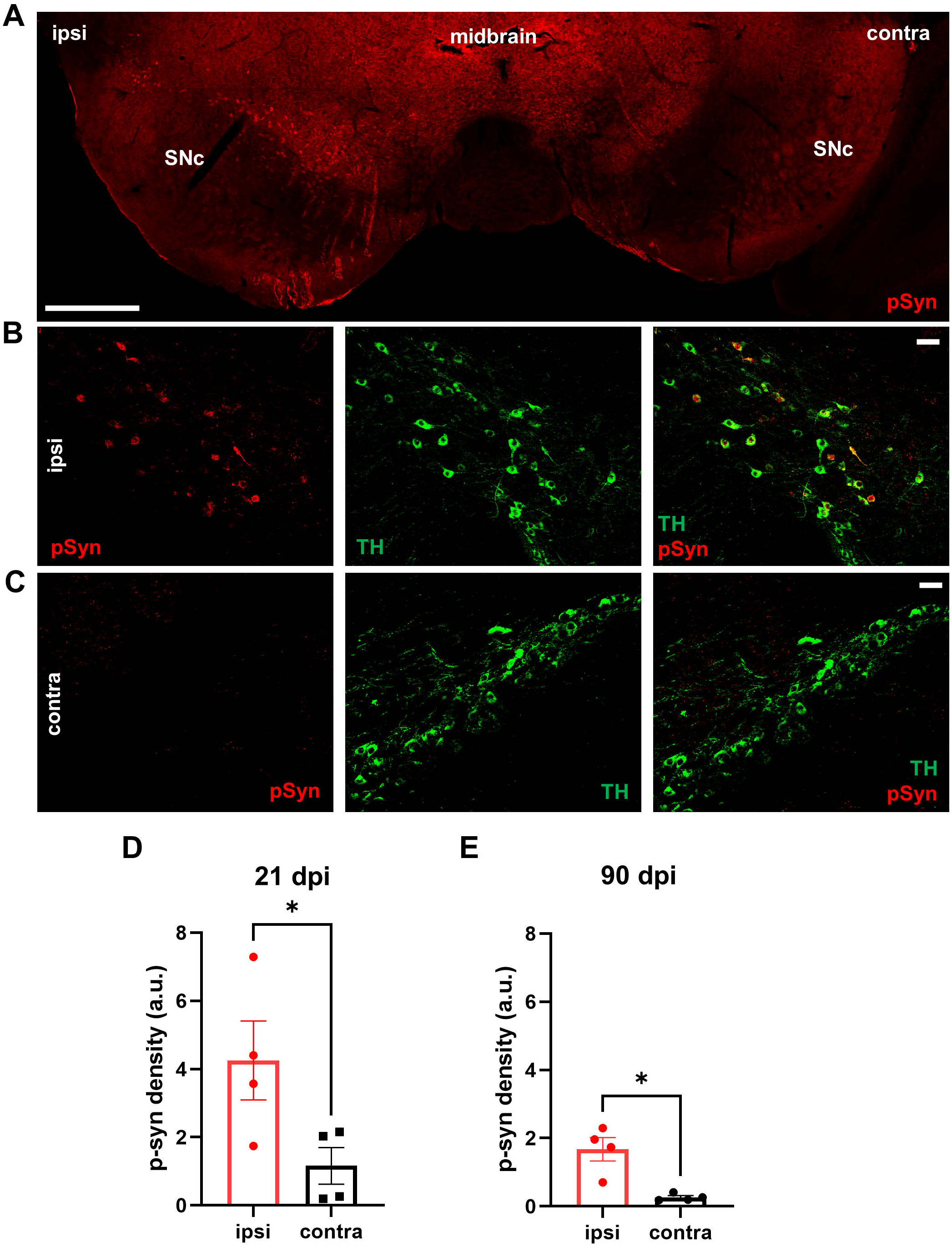
Increased phosphorylation at serine-129 of α-synuclein after hASYN overexpression. **(A)** Detection of α-synuclein phosphorylated at serine 129 (pSyn; red) in a midbrain section from a DAT^Cre^ mouse unilaterally injected with AAV-DIO-hASYN^WT^ 21 dpi; scale bar: 500 µm. **(B-C)** Higher resolution confocal images from SNc stained for pSyn (red) and TH (green) from the ipsilateral **(B)** or contralateral side **(C)**; scale bars: 50 µm. **(D-E)** Densitometric quantification of pSyn levels in midbrain DA neurons of ipsi- and contralateral sides at 21 **(D)** and 90 **(E)** days post virus infusion; n=4 mice/group; two-tailed paired Student’s t-test; 21dpi: t=3.756, df=3, *p=0.0330; 90 dpi: t=4.378, df=3, *p=0.0221.

## Discussion

We have generated a novel viral vector for cell-type specific overexpression of wildtype human α-synuclein in genetically-defined neuronal populations based on the expression of Cre recombinase. In this study we have focused on cell-autonomous effects after human wildtype α-synuclein overexpression in mouse DA neurons within the SNc, the region most severely degenerating in PD.

We found that three weeks after injection of AAV-DIO-hASYN^WT^ into the SNc of DAT^Cre^ mice, human α-synuclein expression within transduced DA neurons was significantly elevated compared to endogenous murine α-synuclein at both the level of cell bodies in the SNc and in striatal terminals. However, despite abundant overexpression in SNc DA neurons and hyperphosphorylation at serine 129, α-synuclein did not grossly disrupt the integrity of DA terminals or neuron numbers up to 90 days after injection.

These findings were unexpected since many publications indicated degeneration of DA neurons in the ventral midbrain after virus-mediated overexpression of α-synuclein in rat, mouse and monkey within just a few weeks [17, 21, 22, 46]. However, there are also several major differences between these studies and ours that may explain the apparent discrepancies: The first and most obvious difference is that we restricted α-synuclein overexpression to DA neurons within the SNc while the other studies relied on non-cell type specific α-synuclein vectors. To our knowledge, there are only two other studies that relied on cell-type specific AAVs to overexpress α-synuclein so far: a study by Grames et al. showed that overexpression of wildtype human α-synuclein (using AAV_9_-EF1α-DIO-hASYN^WT^) in the SNc of TH^Cre^ rats did not induce any loss of TH neurons four weeks after injection [14], which is in line with our findings in mice. And we showed previously that overexpression of the disease-associated variant α-synuclein^A53T^ (using AAV_DJ_-hSyn1-DIO-hASYN^A53T^) did not induce significant changes in the number of TH mRNA^+^ neurons in the SNc three months after virus injection in DAT^Cre^ mice [43]. Hence, cell-type specific vs. non-specific vectors to express α-synuclein in SNc may be essential regarding the observed differences in neuropathology. It remains unclear whether cell-autonomous effects within DA neurons of the SNc make these cells particularly vulnerable to α-synuclein-driven degeneration, or whether α-synuclein in other neurons or glia drive α-synuclein pathology through neural circuit connectivity, inflammatory signals from microglia or immune cells, or whether a complex interaction of multiple processes is required to initiate neurodegeneration. Accordingly, the interaction of different cell types overexpressing α-synuclein rather than overexpression in just one cell type may explain why degeneration was not evident with our vector, at least up to 90 days post injection. Interestingly, lack of overt neurodegeneration in dopamine cell bodies or terminals was also observed in a transgenic mouse line that overexpressed wildtype human α-synuclein in DA neurons driven by the rat TH promoter even at old age [36, 39].

*II)* We used the human Synapsin 1 (hSyn1) promoter to drive α-synuclein expression, while most other commonly used AAVs for α-synuclein overexpression used very strong promoters such as CBA (ß-actin promoter with enhancer elements from the CMV promoter) or CMV. Use of the latter promoters may lead to even higher α-synuclein expression levels than with the hSyn1 or EF1a promoter [14, 43] and therefore they may be more likely to induce toxicity. Note that there are also reports showing that ‘inert’ proteins like GFP can cause toxicity suggesting that overexpression of any protein may become toxic once a critical threshold is reached [4, 23]. Finally, it is important to mention that there may be heterogeneity in terms of actual protein expression levels within different SNc dopamine neurons: some cells could express many fold higher levels of α-synuclein whereas other cells may express only a bit more compared to a non-overexpressing cell. Therefore, the individual expression levels are not readily assessable.

But is elevating the levels of α-synuclein *per se* sufficient to induce degeneration? The genetic findings that SNCA multiplication causes familial PD imply that too much α-synuclein may be toxic [5, 41], however, whether toxicity results from gain-of-toxic-function or loss-of-function remains controversial. For instance, there are studies that suggest better motor and cognitive outcomes in patients with SNCA multiplication compared to PD patients with ‘normal’ SNCA expression [30] implying that elevation of α-synuclein may exert beneficial effects during ongoing neurodegeneration. Interestingly, earlier animal work on α-synuclein suggested that elevated levels of α-synuclein may not be toxic but instead could represent a compensatory and protective cellular response to neuronal injury [20].

Our experiments indicate that the simple elevation of α-synuclein was not sufficient to induce DA neuron toxicity within the three months observation period. It may be that mice are intrinsically more resistant to this type of neuronal stress, longer incubation times are needed or that a ‘second hit’ such as additional oxidative stress or α-synuclein ‘seeds’ are required to induce degeneration.

While we observed no signs of gross DA neurodegeneration after α-synuclein overexpression in SNc neurons, mice clearly exhibited DA dysfunction: mice unilaterally overexpressing α-synuclein displayed increased spontaneous motor activity in the open-field at 21 and 90 days after virus injection; and this increased motor activity was accompanied by elevated striatal DA tissue levels. Since most of the tissue DA is contained in synaptic vesicles [9, 26] it suggests that overexpression of α-synuclein may have affected vesicular DA uptake, vesicular storage, vesicle numbers and/or vesicle turnover. Interestingly, mice lacking α-synuclein display reduced striatal dopamine tissue content further implicating α-synuclein in the dopamine vesicle cycle [1].

In fact, α-synuclein exerts complex effects on vesicular exocytosis and transmitter release. For instance, α-synuclein knockout mice display increased dopamine release in response to paired stimuli [1], while elevated levels of α-synuclein have been largely proposed to decrease the vesicular release of neurotransmitters through effects on the synaptic vesicle cycle [28, 33]. In line with this, several studies have found reduced DA release after overexpression of α-synuclein prior to degeneration using α-synuclein AAV infusions in rats or BAC transgenic SNCA overexpressing mice [11, 19]. Yet, other studies indicate elevated extracellular DA levels in a Thy1-α-synuclein transgenic mouse indicating increased tonic release [27]; or increased vesicular DA release in young but not old adult PDGF-human α-synuclein mice [31]. Together, these findings underline complex effects of α-synuclein on the synaptic vesicle cycle.

In line with other AAV-mediated α-synuclein overexpression studies we have also observed increased phosphorylation of α-synuclein at serine-129 at 21 and 90 days after virus injection. While serine-129 phosphorylation is typically considered to be a pathological feature of synucleinopathies [2, 10], more recent studies suggest that it is also augmented in response to neuronal activity [35, 38]. Hence, the increase in serine-129 phosphorylation after overexpression of α-synuclein in DA neurons does not seem to indicate ongoing neurodegeneration, at least during our observation period. Rather, it could indicate that DA neuron activity may have increased in response to elevated α-synuclein.

### Conclusion

Together, our data suggest that elevated levels of wildtype human α-synuclein are not acutely toxic to DA neurons but that increased α-synuclein results in augmented DA tissue levels and hyperactivity. The lack of α-synuclein overexpression toxicity also makes our novel cell-type specific AAV a great tool to further investigate the physiological role of α-synuclein within defined neuronal populations.

## Declarations

### Ethics approval and consent to participate

All mice were used in accordance with protocols approved by the Animal Welfare Committee of the Medical University of Vienna and the Austrian Federal Ministry of Science and Research (BMBWF licenses 2021-0.373.073 and 2023-0.515.074) or protocols approved by the University of California, San Diego (UCSD) Institutional Animal Care and Use Committee.

## Consent for publication

Not applicable.

## Availability of data and material

The datasets used and/or analysed during the current study are available from the corresponding author on reasonable request.

## Competing interests

The authors declare that they have no competing interests.

## Funding

This work was supported by the Austrian Science Fund (FWF P-35871, FWF-P-36125 and FWF-P-35719), NIH-K99-AG059834 and a BBRF NARSAD Young Investigator award 30784 (to TS) and joint efforts of the Michael J. Fox Foundation for Parkinson’s Research (MJFF) and the Aligning Science Across Parkinson’s (ASAP) initiative. MJFF administers the grant (ASAP-020600) on behalf of ASAP and itself (to TSH).

## Authors’ Contributions

TS designed research. SGM, FL, IL, CG and TS performed research. CP helped with HPLC setup and analysis. TSH contributed with infrastructural support, data evaluation and discussion. TS wrote the manuscript with input from all authors. All authors read and approved the manuscript.

## Acknowledgements

We thank Brian Spencer and Robert Rissman (UCSD) for pLV-α-synuclein plasmids.

---

**Supplemental Figure 1: Expression of AAV-DIO-hASYN is specific to Cre-expressing mouse lines. (A)** Injection of AAV-DIO-hASYN^WT^ into the left SNc of DAT^Cre^ or DAT^+/+^ (C57BL/6) mice shows that hASYN is only expressed in the presence of Cre recombinase (left) but not in its absence. **(B)** Expression of AAV-DIO-hASYN^WT^ in striatal cholinergic interneurons of ChAT^Cre^ mice. **(C, D)** Expression of AAV-DIO-hASYN^WT^ in the striatal direct pathway of D1^Cre^ **(C)** and in the striatal indirect pathway of A2a^Cre^ **(D)** mice. Co-staining was performed with specific antibodies directed against D1 **(C)** or D2 **(D)** receptors. CPu = caudate/putamen, GPe = globus pallidus *externus*, GPi = globus pallidus *internus*, SNr = substantia nigra *pars reticulata*.

**Supplemental Figure 2: Uncropped immunoblots used for striatal alpha-synuclein detection and quantification.** Upper panels: Immunoblots of striatal protein lysates probed for human alpha synuclein (15G7; green), total alpha-synuclein (Syn1; red) and loading control (GAPDH; blue) n=4 mice/group. L = protein standard ladder. Lower panel: Ponceau S staining of nitrocellulose membrane after wet transfer.

**Supplemental Figure 3: Molar ratios of DOPAC/DA and HVA/DA.**

## List of abbreviations

AAV: adeno-associated virus
ChAT: choline acetyltransferase
CMV: cytomegalovirus promoter
CPu: caudate/putamen
DA: dopamine
DAT^iCre^: mice expressing Cre recombinase driven by dopamine transporter regulatory elements
DIO: double inverted open reading frame
DOPAC: 3,4-dihydroxyphenylacetic acid
DLB: dementia with Lewy bodies
GPe: globus pallidus *externus*
GPi: globus pallidus *internus*
hASYN: human alpha-synuclein
hSyn1: human synapsin 1 promoter
HPLC: high performance liquid chromatography
HVA: homovanillic acid
LB: Lewy bodies
SNc: substantia nigra *pars compacta*
SNCA: alpha-synuclein gene
SNr: substantia nigra *pars reticulata*
TH: tyrosine hydroxylase
VMAT2: vesicular monoamine transporter 2

## Notes

### Competing Interest Statement

The authors have declared no competing interest.

## References

1 Abeliovich A, Schmitz Y, Farinas I, Choi-Lundberg D, Ho WH, Castillo PE, Shinsky N, Verdugo JM, Armanini M, Ryan Aet al (2000) Mice lacking alpha-synuclein display functional deficits in the nigrostriatal dopamine system. Neuron 25: 239–252 Doi 10.1016/s0896-6273(00)80886-7

2 Albin RL, Young AB, Penney JB, Handelin B, Balfour R, Anderson KD, Markel DS, Tourtellotte WW, Reiner A (1990) Abnormalities of striatal projection neurons and N-methyl-D-aspartate receptors in presymptomatic Huntington’s disease. N Engl J Med 322: 1293–1298 Doi 10.1056/NEJM199005033221807

3. Anderson JP, Walker DE, Goldstein JM, de Laat R, Banducci K, Caccavello RJ, Barbour R, Huang J, Kling K, Lee M et al (2006) Phosphorylation of Ser-129 is the dominant pathological modification of alpha-synuclein in familial and sporadic Lewy body disease. J Biol Chem 281: 29739–29752 Doi 10.1074/jbc.M600933200

4. Buck SA, Steinkellner T, Aslanoglou D, Villeneuve M, Bhatte SH, Childers VC, Rubin SA, De Miranda BR, O’Leary EI, Neureiter EG et al (2021) Vesicular glutamate transporter modulates sex differences in dopamine neuron vulnerability to age-related neurodegeneration. Aging Cell 20: e13365 Doi 10.1111/acel.13365

5 Chartier-Harlin MC, Kachergus J, Roumier C, Mouroux V, Douay X, Lincoln S, Levecque C, Larvor L, Andrieux J, Hulihan Met al (2004) Alpha-synuclein locus duplication as a cause of familial Parkinson’s disease. Lancet 364: 1167–1169 Doi 10.1016/S0140-6736(04)17103-1

6 Chesselet MF, Richter F (2011) Modelling of Parkinson’s disease in mice. Lancet Neurol 10: 1108–1118 Doi 10.1016/S1474-4422(11)70227-7

7 Cookson MR (2009) alpha-Synuclein and neuronal cell death. Mol Neurodegener 4: 9 Doi 10.1186/1750-1326-4-9

8 Damier P, Hirsch EC, Agid Y, Graybiel AM (1999) The substantia nigra of the human brain. II. Patterns of loss of dopamine-containing neurons in Parkinson’s disease. Brain 122 (Pt 8): 1437–1448 Doi 10.1093/brain/122.8.1437

9 Fon EA, Pothos EN, Sun BC, Killeen N, Sulzer D, Edwards RH (1997) Vesicular transport regulates monoamine storage and release but is not essential for amphetamine action. Neuron 19: 1271–1283 Doi 10.1016/s0896-6273(00)80418-3

10 Fujiwara H, Hasegawa M, Dohmae N, Kawashima A, Masliah E, Goldberg MS, Shen J, Takio K, Iwatsubo T (2002) alpha-Synuclein is phosphorylated in synucleinopathy lesions. Nat Cell Biol 4: 160–164 Doi 10.1038/ncb748

11 Gaugler MN, Genc O, Bobela W, Mohanna S, Ardah MT, El-Agnaf OM, Cantoni M, Bensadoun JC, Schneggenburger R, Knott GW et al (2012) Nigrostriatal overabundance of alpha-synuclein leads to decreased vesicle density and deficits in dopamine release that correlate with reduced motor activity. Acta Neuropathol 123: 653–669 Doi 10.1007/s00401-012-0963-y

12 Gibb WR, Lees AJ (1988) The relevance of the Lewy body to the pathogenesis of idiopathic Parkinson’s disease. J Neurol Neurosurg Psychiatry 51: 745–752 Doi 10.1136/jnnp.51.6.745

13 Giguere N, Burke Nanni S, Trudeau LE (2018) On Cell Loss and Selective Vulnerability of Neuronal Populations in Parkinson’s Disease. Front Neurol 9: 455 Doi 10.3389/fneur.2018.00455

14 Grames MS, Dayton RD, Jackson KL, Richard AD, Lu X, Klein RL (2018) Cre-dependent AAV vectors for highly targeted expression of disease-related proteins and neurodegeneration in the substantia nigra. FASEB J 32: 4420–4427 Doi 10.1096/fj.201701529RR

15 Heusner CL, Beutler LR, Houser CR, Palmiter RD (2008) Deletion of GAD67 in dopamine receptor-1 expressing cells causes specific motor deficits. Genesis 46: 357–367 Doi 10.1002/dvg.20405

16 Hirsch EC, Graybiel AM, Agid Y (1989) Selective vulnerability of pigmented dopaminergic neurons in Parkinson’s disease. Acta Neurol Scand Suppl 126: 19–22 Doi 10.1111/j.1600-0404.1989.tb01778.x

17 Ip CW, Klaus LC, Karikari AA, Visanji NP, Brotchie JM, Lang AE, Volkmann J, Koprich JB (2017) AAV1/2-induced overexpression of A53T-alpha-synuclein in the substantia nigra results in degeneration of the nigrostriatal system with Lewy-like pathology and motor impairment: a new mouse model for Parkinson’s disease. Acta Neuropathol Commun 5: 11 Doi 10.1186/s40478-017-0416-x

18. Iwai A, Masliah E, Yoshimoto M, Ge N, Flanagan L, de Silva HA, Kittel A, Saitoh T (1995) The precursor protein of non-A beta component of Alzheimer’s disease amyloid is a presynaptic protein of the central nervous system. Neuron 14: 467–475 Doi 10.1016/0896-6273(95)90302-x

19 Janezic S, Threlfell S, Dodson PD, Dowie MJ, Taylor TN, Potgieter D, Parkkinen L, Senior SL, Anwar S, Ryan Bet al (2013) Deficits in dopaminergic transmission precede neuron loss and dysfunction in a new Parkinson model. Proc Natl Acad Sci U S A 110: E4016–4025 Doi 10.1073/pnas.1309143110

20 Kholodilov NG, Neystat M, Oo TF, Lo SE, Larsen KE, Sulzer D, Burke RE (1999) Increased expression of rat synuclein in the substantia nigra pars compacta identified by mRNA differential display in a model of developmental target injury. J Neurochem 73: 2586–2599 Doi 10.1046/j.1471-4159.1999.0732586.x

21 Kirik D, Annett LE, Burger C, Muzyczka N, Mandel RJ, Bjorklund A (2003) Nigrostriatal alpha-synucleinopathy induced by viral vector-mediated overexpression of human alpha-synuclein: a new primate model of Parkinson’s disease. Proc Natl Acad Sci U S A 100: 2884–2889 Doi 10.1073/pnas.0536383100

22 Kirik D, Rosenblad C, Burger C, Lundberg C, Johansen TE, Muzyczka N, Mandel RJ, Bjorklund A (2002) Parkinson-like neurodegeneration induced by targeted overexpression of alpha-synuclein in the nigrostriatal system. J Neurosci 22: 2780–2791 Doi 10.1523/JNEUROSCI.22-07-02780.2002

23 Klein RL, Dayton RD, Leidenheimer NJ, Jansen K, Golde TE, Zweig RM (2006) Efficient neuronal gene transfer with AAV8 leads to neurotoxic levels of tau or green fluorescent proteins. Mol Ther 13: 517–527 Doi 10.1016/j.ymthe.2005.10.008

24 Klein RL, King MA, Hamby ME, Meyer EM (2002) Dopaminergic cell loss induced by human A30P alpha-synuclein gene transfer to the rat substantia nigra. Hum Gene Ther 13: 605–612 Doi 10.1089/10430340252837206

25 Kruger R, Kuhn W, Muller T, Woitalla D, Graeber M, Kosel S, Przuntek H, Epplen JT, Schols L, Riess O (1998) Ala30Pro mutation in the gene encoding alpha-synuclein in Parkinson’s disease. Nat Genet 18: 106–108 Doi 10.1038/ng0298-106

26 Kuczenski R (1977) Differential effects of reserpine and tetrabenazine on rat striatal synaptosomal dopamine biosynthesis and synaptosomal dopamine pools. J Pharmacol Exp Ther 201: 357–367

27 Lam HA, Wu N, Cely I, Kelly RL, Hean S, Richter F, Magen I, Cepeda C, Ackerson LC, Walwyn Wet al (2011) Elevated tonic extracellular dopamine concentration and altered dopamine modulation of synaptic activity precede dopamine loss in the striatum of mice overexpressing human alpha-synuclein. J Neurosci Res 89: 1091–1102 Doi 10.1002/jnr.22611

28 Larsen KE, Schmitz Y, Troyer MD, Mosharov E, Dietrich P, Quazi AZ, Savalle M, Nemani V, Chaudhry FA, Edwards RH et al (2006) Alpha-synuclein overexpression in PC12 and chromaffin cells impairs catecholamine release by interfering with a late step in exocytosis. J Neurosci 26: 11915–11922 Doi 10.1523/JNEUROSCI.3821-06.2006

29 Lo Bianco C, Ridet JL, Schneider BL, Deglon N, Aebischer P (2002) alpha - Synucleinopathy and selective dopaminergic neuron loss in a rat lentiviral-based model of Parkinson’s disease. Proc Natl Acad Sci U S A 99: 10813–10818 Doi 10.1073/pnas.152339799

30 Markopoulou K, Biernacka JM, Armasu SM, Anderson KJ, Ahlskog JE, Chase BA, Chung SJ, Cunningham JM, Farrer M, Frigerio Ret al (2014) Does alpha-synuclein have a dual and opposing effect in preclinical vs. clinical Parkinson’s disease? Parkinsonism Relat Disord 20: 584–589; discussion 584 Doi 10.1016/j.parkreldis.2014.02.021

31 Medina-Luque J, Piechocinski P, Feyen P, Sgobio C, Herms J (2024) Striatal dopamine neurotransmission is altered in age-and region-specific manner in a Parkinson’s disease transgenic mouse. Sci Rep 14: 164 Doi 10.1038/s41598-023-49600-5

32 Murphy DD, Rueter SM, Trojanowski JQ, Lee VM (2000) Synucleins are developmentally expressed, and alpha-synuclein regulates the size of the presynaptic vesicular pool in primary hippocampal neurons. J Neurosci 20: 3214–3220 Doi 10.1523/JNEUROSCI.20-09-03214.2000

33 Nemani VM, Lu W, Berge V, Nakamura K, Onoa B, Lee MK, Chaudhry FA, Nicoll RA, Edwards RH (2010) Increased expression of alpha-synuclein reduces neurotransmitter release by inhibiting synaptic vesicle reclustering after endocytosis. Neuron 65: 66–79 Doi 10.1016/j.neuron.2009.12.023

34 Parkkinen L, O’Sullivan SS, Collins C, Petrie A, Holton JL, Revesz T, Lees AJ (2011) Disentangling the relationship between lewy bodies and nigral neuronal loss in Parkinson’s disease. J Parkinsons Dis 1: 277–286 Doi 10.3233/JPD-2011-11046

35 Parra-Rivas LA, Madhivanan K, Aulston BD, Wang L, Prakashchand DD, Boyer NP, Saia-Cereda VM, Branes-Guerrero K, Pizzo DP, Bagchi Pet al (2023) Serine-129 phosphorylation of alpha-synuclein is an activity-dependent trigger for physiologic protein-protein interactions and synaptic function. Neuron 111: 4006–4023 e4010 Doi 10.1016/j.neuron.2023.11.020

36 Peneder TM, Scholze P, Berger ML, Reither H, Heinze G, Bertl J, Bauer J, Richfield EK, Hornykiewicz O, Pifl C (2011) Chronic exposure to manganese decreases striatal dopamine turnover in human alpha-synuclein transgenic mice. Neuroscience 180: 280–292 Doi 10.1016/j.neuroscience.2011.02.017

37 Polymeropoulos MH, Lavedan C, Leroy E, Ide SE, Dehejia A, Dutra A, Pike B, Root H, Rubenstein J, Boyer Ret al (1997) Mutation in the alpha-synuclein gene identified in families with Parkinson’s disease. Science 276: 2045–2047 Doi 10.1126/science.276.5321.2045

38 Ramalingam N, Jin SX, Moors TE, Fonseca-Ornelas L, Shimanaka K, Lei S, Cam HP, Watson AH, Brontesi L, Ding Let al (2023) Dynamic physiological alpha-synuclein S129 phosphorylation is driven by neuronal activity. NPJ Parkinsons Dis 9: 4 Doi 10.1038/s41531-023-00444-w

39. Richfield EK, Thiruchelvam MJ, Cory-Slechta DA, Wuertzer C, Gainetdinov RR, Caron MG, Di Monte DA, Federoff HJ (2002) Behavioral and neurochemical effects of wild-type and mutated human alpha-synuclein in transgenic mice. Exp Neurol 175: 35–48 Doi 10.1006/exnr.2002.7882

40 Saunders A, Johnson CA, Sabatini BL (2012) Novel recombinant adeno-associated viruses for Cre activated and inactivated transgene expression in neurons. Front Neural Circuits 6: 47 Doi 10.3389/fncir.2012.00047

41 Singleton AB, Farrer M, Johnson J, Singleton A, Hague S, Kachergus J, Hulihan M, Peuralinna T, Dutra A, Nussbaum Ret al (2003) alpha-Synuclein locus triplication causes Parkinson’s disease. Science 302: 841 Doi 10.1126/science.1090278

42 Spillantini MG, Schmidt ML, Lee VM, Trojanowski JQ, Jakes R, Goedert M (1997) Alpha-synuclein in Lewy bodies. Nature 388: 839–840 Doi 10.1038/42166

43 Steinkellner T, Conrad WS, Kovacs I, Rissman RA, Lee EB, Trojanowski JQ, Freyberg Z, Roy S, Luk KC, Lee VM et al (2022) Dopamine neurons exhibit emergent glutamatergic identity in Parkinson’s disease. Brain 145: 879–886 Doi 10.1093/brain/awab373

44 Steinkellner T, Zell V, Farino ZJ, Sonders MS, Villeneuve M, Freyberg RJ, Przedborski S, Lu W, Freyberg Z, Hnasko TS (2018) Role for VGLUT2 in selective vulnerability of midbrain dopamine neurons. J Clin Invest 128: 774–788 Doi 10.1172/JCI95795

45 Taguchi K, Watanabe Y, Tsujimura A, Tanaka M (2016) Brain region-dependent differential expression of alpha-synuclein. J Comp Neurol 524: 1236–1258 Doi 10.1002/cne.23901

46 Theodore S, Cao S, McLean PJ, Standaert DG (2008) Targeted overexpression of human alpha-synuclein triggers microglial activation and an adaptive immune response in a mouse model of Parkinson disease. J Neuropathol Exp Neurol 67: 1149–1158 Doi 10.1097/NEN.0b013e31818e5e99

47 Zarranz JJ, Alegre J, Gomez-Esteban JC, Lezcano E, Ros R, Ampuero I, Vidal L, Hoenicka J, Rodriguez O, Atares Bet al (2004) The new mutation, E46K, of alpha-synuclein causes Parkinson and Lewy body dementia. Ann Neurol 55: 164-173 Doi 10.1002/ana.10795

