## Supplementary figures and images for "Viral overexpression of human alpha-synuclein in mouse substantia nigra dopamine neurons results in hyperdopaminergia but no neurodegeneration"

### Supplemental Figure 1

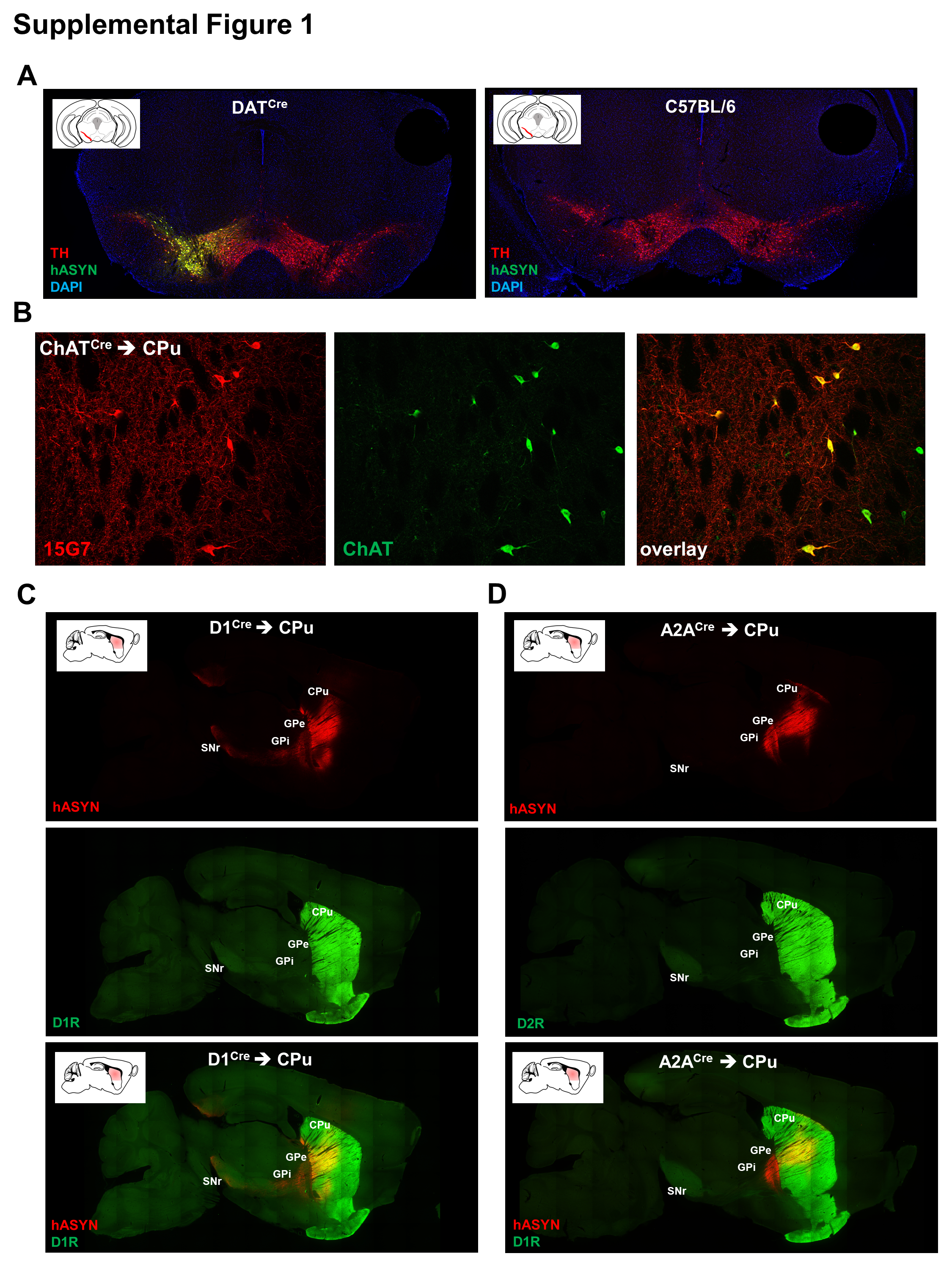

### Supplemental Figure 2

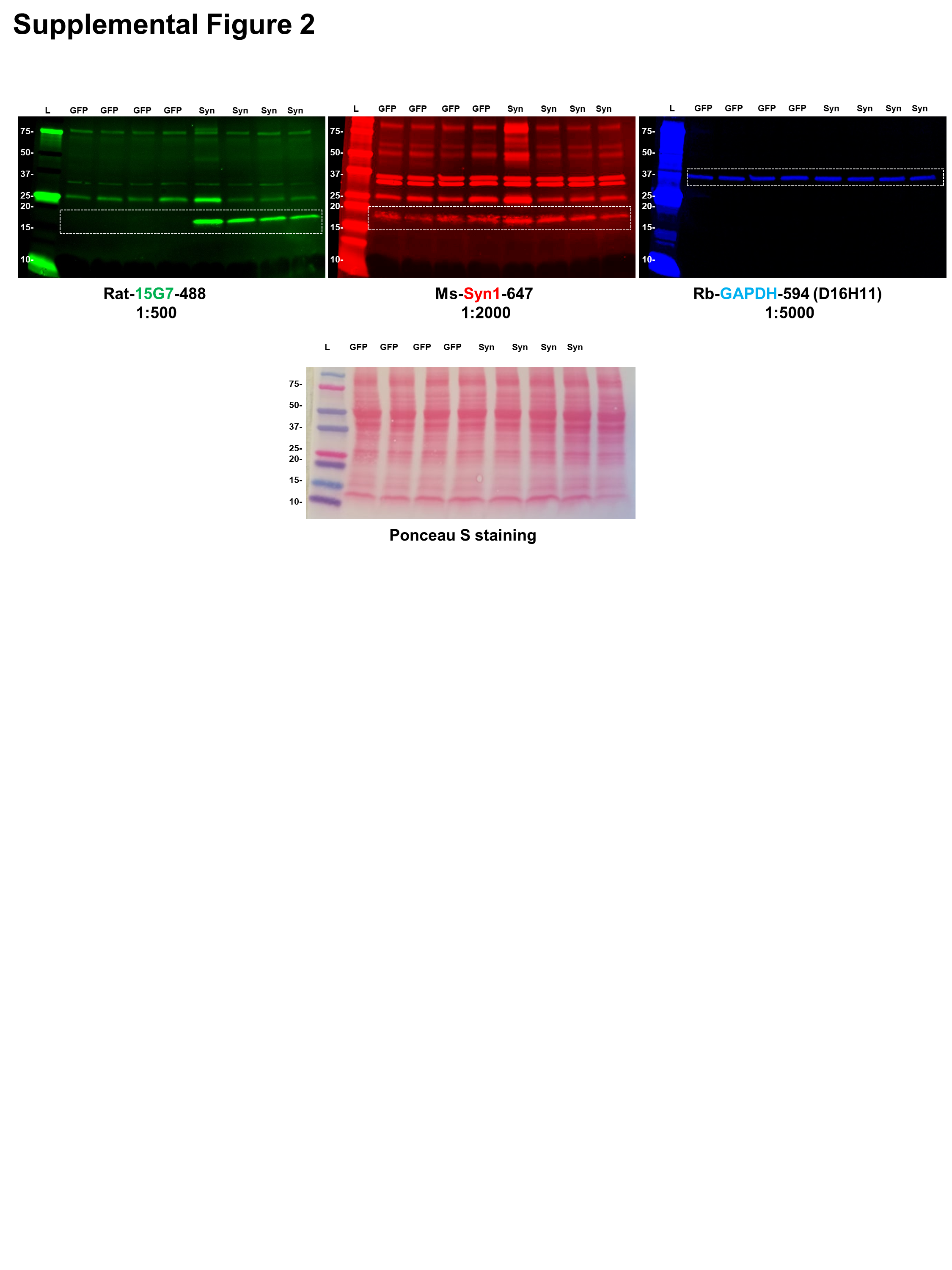

### Supplemental Figure 3

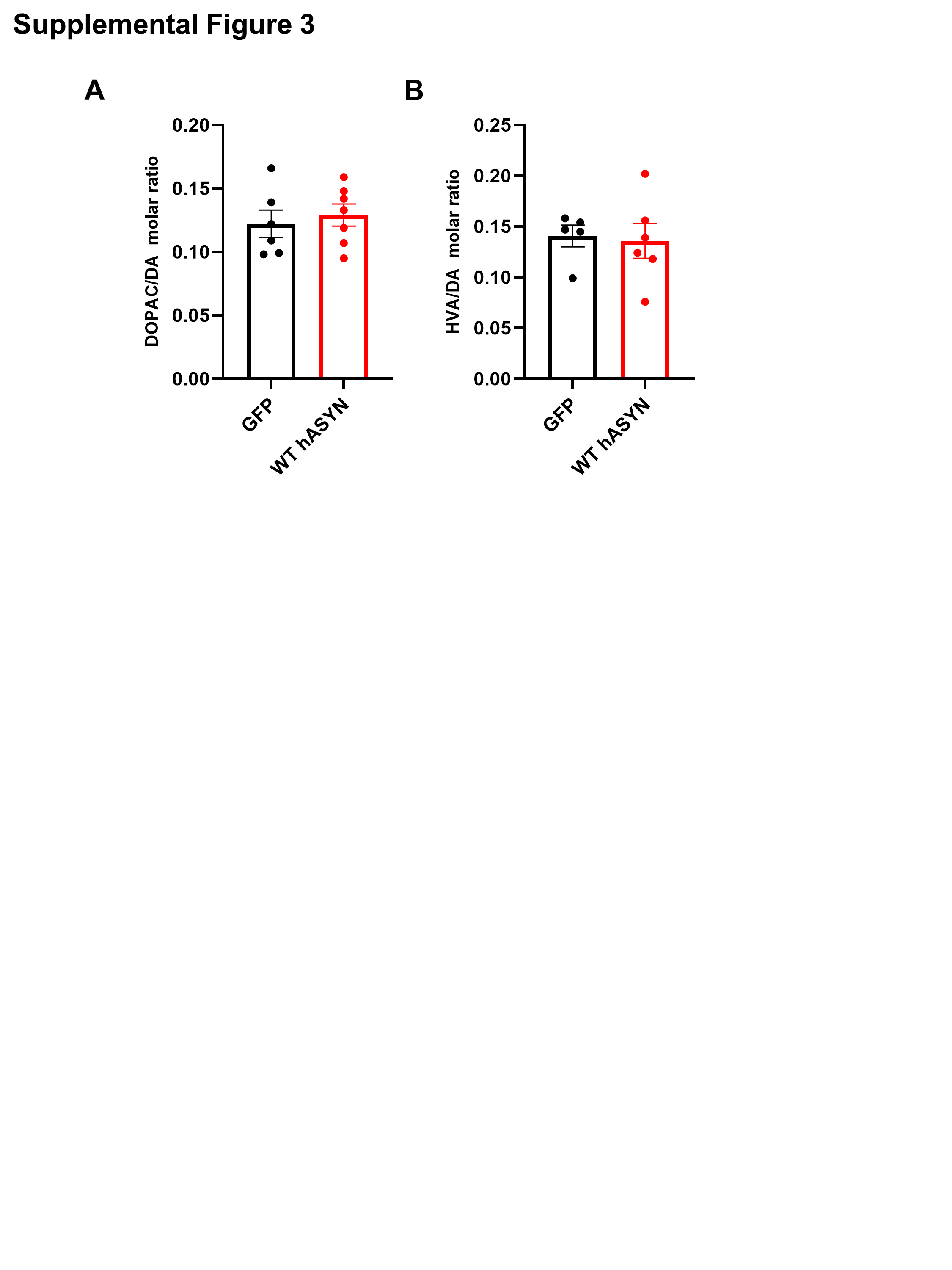
